# The High-quality genome assembly of *Lactarius hatsudake*

**DOI:** 10.1101/2022.08.15.504028

**Authors:** Airong Shen, Chen Luo, Yun Tan, Baoming Shen, Lina Liu, Liangbin Zeng, Zhuming Tan, Jilie Li

**Affiliations:** Central South University of Forestry and Technology, Changsha 410004, China; Hunan Academy of Forestry, Changsha, 410004, China; Institute of Bast Fiber Crops, Chinese Academy of Agricultural Sciences, Changsha 410205, China

**Author notes:** Corresponding authors: Liangbin Zeng, Institute of Bast Fiber Crops, Chinese Academy of Agricultural Sciences, No 348 Xianjiahu West Road, Changsha, Hunan, 410205, China,. And Zhuming Tan, Hunan Academy of Forestry, No. 658 Shaoshan South Road, Changsha, 410004, China,. And Jilie Li, Central South University of Forestry and Technology, No. 498 Shaoshan South Road, Changsha, Hunan, 410004, China,. Airong Shen and Chen Luo contributed equally to this work.

**Keywords:** *Lactarius hatsudake*, Illumina, PacBio, Genome

## Abstract

*Lactarius hatsudake* is a species of *Lactarius* commonly found in pine forests, is edible with a delicious and nutritious fruiting body and exhibits medicinal properties. It is an ideal natural multi-functional food with bioactive components including fungal polysaccharides, crude fiber, unsaturated fatty acids, nucleic acid derivatives, various amino acids, and vitamins. However, biological and genomic analyses of this mycorrhizal mushroom are sparse, thereby hindering large-scale cultivation. Previously, we isolated and screened *L. hatsudake* JH5 strains and have applied our garnered knowledge to the large-scale cultivation of mycorrhizal seedlings. In this study, we produced a high-quality genome assembly of L. hatsudake JH5 by combining Illumina paired-end (PE) and PacBio single molecule real-time (SMRT) sequencing, resulting in PacBio SMRT reads of 7.67 Gb and Illumina Pair-End reads of 1560 Mb. Based on the distribution of k-mer frequencies, the genome size of this strain was estimated to be 63.84 Mb (1.14% heterozygosity). Based on *de novo* genome assembly, the final genome size was determined to be 76.7 Mb, with scaffold N50 of 223.2 kb and N90 of 54.5 kb, and a GC content of 54.38%. BUSCO assessment showed that genome completeness was 89.0%. The N50 length of the JH5 genome was 43.6% longer than that of the previously published *L. hatsudake* MG20 genome. This high-quality *L. hatsudake* genome assembly will facilitate research on the functional genome, molecular breeding, yield enhancement and sustainability of *L. hatsudake* cultivation.

---

*Lactarius hatsudake* Tanaka, also known as red milk mushroom, is a high-quality wild edible and medicinal mycorrhizal fungus that is symbiotic with Pinaceaeor and Lophiraceae. It is principally distributed in North America, Europe, and Southeast Asia within China, Korea, Thailand, and Japan (Fang et al. 2006). The fruiting bodies of *L. hatsudake* are nutritious and rich in bioactive components such as fungal polysaccharides, crude fiber, unsaturated fatty acids, nucleic acid derivatives, various amino acids and vitamins (Miyazawa et al. 2010; Wang Li et al. 2016). Previous studies have shown that *L. hatsudake* is able to alleviate symptoms of Diabetes patients, improve immune responses, and inhibit pathogenic bacteria (Tako et al. 2012; Zhang et al. 2007), thus serving as an ideal natural multi-functional high-grade food source. *L. hatsudake* is also known as cold fungus, wild goose fungus, gong fungus, silk fungus, and purple flower fungus and has become a major species in the wild edible mushroom trade in southeastern China (Li et al. 2020) with popularity extending to Japan, South Korea, Thailand, among other locations (Miyazawa et al. 2010; Tako et al. 2012). The mushroom can be consumed fresh, frozen, or processed into mushroom oil.

Successful cultivation of *L. hatsudake* has been a long-standing desire within production regions. Initially, *L. hatsudake*–*Pinus massoniana* Lamb was obtained by tissue isolation and mycorrhizal synthesis techniques (Tan 2005; Tan 2006; Tan et al. 2007). Currently, the red milk mushroom plantation in Hunan Province covers 118 hectares with the fruiting bodies produced in 3-4 years after mycorrhizal seeding. Successful cultivation of *L. hatsudake* has led to a novel forest economy with associated ecological and economic benefits. However, due to the extended cultivation time and a need for extensive plantation management, it has been difficult to reliably achieve stable and high yields. A lack of comprehensive whole genome analysis for *L. hatsudake* has also served to limit mechanistic analysis. What is currently known is that, based on whole-genome sequencing, sexual reproduction was found to be heterotypical coordinated in *Tuber melanosporum* (Martin et al. 2010), while *Laccaria bicolor* was found to possess a family of genes associated with symbiosis, called the “symbiosis toolbox” (Martin et al. 2008; Martin and Selosse 2008). Studies have revealed a great diversity of genomic landscapes and gene pools within Russulaceae (Lebreton et al. 2022). These results suggest that in-depth analysis of the genomic characteristics of mycorrhizal edible fungi by genome sequencing technology can help to identify and clarify the process of mycorrhiza formation and fruiting body development, thus contributing to an efficient and sustainable cultivation of this edible mycorrhizal fungi.

Deciphering the genome of *L. hatsudake* will provide theoretical support for the process of mycorrhiza and fruiting body formation and may guide further large-scale cultivation. Currently, twenty-four *Lactarius* species have undergone preliminary genomic analysis, with N50 values ranging from 5.0 kb to 261.3 kb (Li et al. 2020, Lebreton et al. (2022). For *L. hatsudake*, using Illumina sequencing technology, Li et al. (2020) reported a draft genome of *L. hatsudake* MG20 with an N50 of 5.0 kb. More recently, using PacBio sequencing platform, Lebreton et al. (2022) published the genome assembly of *L. hatsudake* 109 with an N50 of 261.3 kb.

Previously, we isolated and screened the JH5 strain of *L. hatsudake* from the field, which has been used for large-scale culture of mycorrhizal seedlings. This strain exhibited rapid growth, adaptation to various nutritional environments, high mycorrhizal infection rate, high competitiveness, and genetic stability (Tan et al. 2021). Herein, we performed *de novo* sequencing of JH5 genome by combining the Illumina and PacBio platform, and provided an improved genome assembly of *L. hatsudake*. This study aims to develop foundational genomic resource for gene functional studies as well as molecular breeding of *L. hatsudake* to improve yield.

## Materials and methods

### Fungal strain and DNA extraction

*L. hatsudake* strain JH5 was obtained from the production and promotion base established by the project team in Pumen Village, Jiahe County, Chenzhou City, Hunan Province. *L. hatsudake* JH5 was selected which was stored in the General Microbiology Center of China microbial species Preservation and Administration Committee, registration number CGMCCNo19369. Mycelia of JH5 was grown in 100 ml biotin-aneurine-folic acid (BAF) liquid medium at 22°C, 120 rpm for 14 days in darkness. Then JH5 mycelia were separated from the culture medium, frozen in liquid nitrogen and ground to a fine powder and subjected to genome sequencing. DNA was extracted using the DNA extraction kit from TIANGEN, Beijing, China, and the purity and integrity of genomic DNA was determined by agarose gel electrophoresis.

### Genome sequencing and assembly

The genome of *L. hatsudake* JH5 was *de novo* sequenced using high throughput Illumina Hiseq X-Ten and PacBio RSII long-read sequencing platforms (PacBio P6-C4) at Beijing Novogene Technology. DNA libraries with 350 bp inserts were constructed and sequenced on the Illumina HiseqX-Ten platform. For the PacBio RSII platform, a 20 kb library was generated and sequenced. The genome size of *L. hatsudake* JH5 was estimated by the k-mer method using sequencing data from the Illumina DNA library. A 15-mer frequency distribution analysis of the quality-filtered reads was performed using Jellyfish v2.2.10 (Marcais and Kingsford 2011). Genome size, heterozygosity and repeat content were then estimated by the Genome Scope web tool (Vurture et al. 2017).

The genome of JH5 was *de novo* assembled in three steps. Assembly of contigs was performed with FALCON (version 0.7.0)(Chin et al. 2016). Briefly, the longest 40×reads were selected as “seed” reads for error correction (“pre-assembly”). Pre-assembly in FALCON uses DALigner to perform all-by-all alignments of the raw reads. The FALCON assembly resulted in 312 primary contigs. The initial polishing was performed with Arrow (included in the FALCON-Unzip) exclusively using PacBio (https://www.pacb.com/support/software-downloads/) long reads, and then SSPACE-LongReads was implemented to scaffold the contigs. Finally, Pilon (v 1.23)(Walker et al. 2014) was utilized to further correct the PacBio-corrected contigs with accurate Illumina short reads, and to generate the genome assembly of *L. hatsudake* JH5. The completeness of the JH5 genome assembly was evaluated using BUSCO 3.1.0 (Benchmarking Universal Single-Copy Orthologs) with comparison to lineage dataset fungi_odb9 (Creation date: 2016-10-21, number of species: 85, number of BUSCOs: 290)(Simao et al. 2015).

### Genome annotation

To annotate the assembled JH5 genome, we used funannotate (v1.5.2) (JonLoveetal. 2019) with the pipeline described in https://funannotate.readthedocs.io/en/latest/tutorials.html with the following commands: funannotate mask, to softmask the genome, funannotate training, and funannotate predict to generate preliminary gene models, and consensus gene models [using: AUGUSTUS (Stanke and Waack 2003), GeneMark (Borodovsky and McIninch 1993), and EVidenceModeler (Haas et al. 2008)], and funannotate annotate to add functional annotation, in addition, protein-coding gene models (PCG) models also were identified according to our twelve transcriptome data (PRJNA841037). The rRNA was predicted by using RNAmmer v1.2 (Lagesen t al. 2007), and tRNAs were identified with tRNAscan-SE v1.4 (Lowe and Eddy, 1997). The sRNA was identified by comparing with the Rfam database (Gardner et al. 2009). The functional annotation obtained with funannotate includes Interpro terms, Pfam domains, CAZYmes (CAZY_DB: 201604), secreted proteins, proteases (MEROPS), BUSCO groups, Eggnog annotations, Clusters of Orthologous Groups (COGs), GO ontology, secretion of signal peptides, and transmembrane domains (the full annotation is available in Supplementary Table 4).

## Results and Discussion

### High-quality genome assembly of *L. hatsudake* JH5

To construct a high-quality reference genome of *L. hatsudake* JH5, a total of 7.67 Gb PacBio SMRT reads and 1560 Mb Illumina pair-end reads were generated in this study. The PacBio read lengths ranged from 200 bp to 50,000 bp with an average read length of 7,418 bp (Figure 1A). We estimated the genome size of *L. hatsudake* JH5 as 63.84 Mb with a heterozygosity rate of 1.14% via the distribution of k-mer frequency using Illumina PE reads (Figure 1B). High-quality PacBio SMRT reads were used to assemble the *L. hatsudake* genome. The contigs were then polished using Illumina PE reads,which yielded a draft genome assembly of 76.7 Mb, with contig N50 of 223.2 kb, N90 of 54.5 kb, GC content of 54.4%, and a BUSCO result of 89.0% (Table 1). We annotated 19,616 genes with an average gene length of 1,765 bp. The cumulative length of genes was 34,6 Mb, which accounted for 45.14% of the whole JH5 genome (Table 1,Table 2). The size of contig N50 and BUSCO results were higher than that of the previously published *L. hatsudake* MG20 genome (contig N50:5,268 bp, BUSCO result:84.5%) (Table 1, Table 2). Compared with the N50 obtained only by Illumina sequencing, the N50 length was 44.6 times higher and the number of genes increased by 1.06 times. The Scaffold N(x) length distribution in *L. hatsudake* JH5 was also significantly higher than the length of Scaffold N(x) distribution in *L. hatsudake* MG20 (Figure 2). In total, this *L. hatsudake* genome assembly represents a significant improvement than that of other previously released *Lactarius* genomes (contig N50: 5.0 kb-261.3 kb) (Table 1) (Li et al. 2020,Saier et al. 2014,Lebreton et al. 2022). Compared with *L. hatsudake* 109 genome assembly, JH5 genome show the considerable N50 length (223.2 kb *vs* 261.3 kb), and BUSCO results (89.0% *vs* 87.6*%*). However, the number of scaffolds of JH5 genome was far fewer than that of *L. hatsudake* 109 genome (312 vs 815), which indicated JH5 genome presented here was less fragmented. Furthermore, 2785 additional genes were predicted compared to *L. hatsudake* 109 genome (Table 1, Table 2).

**Table 1.**
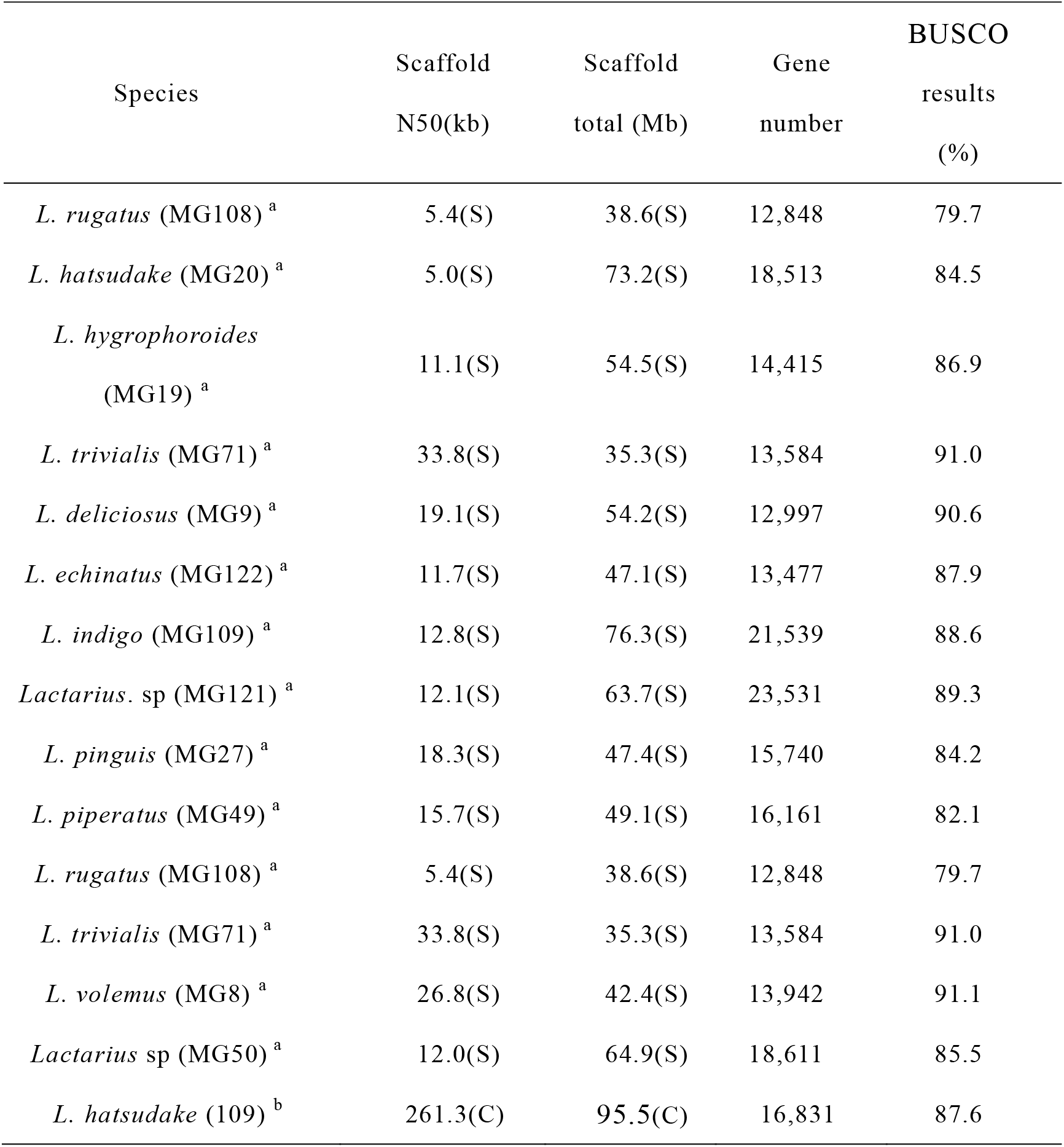
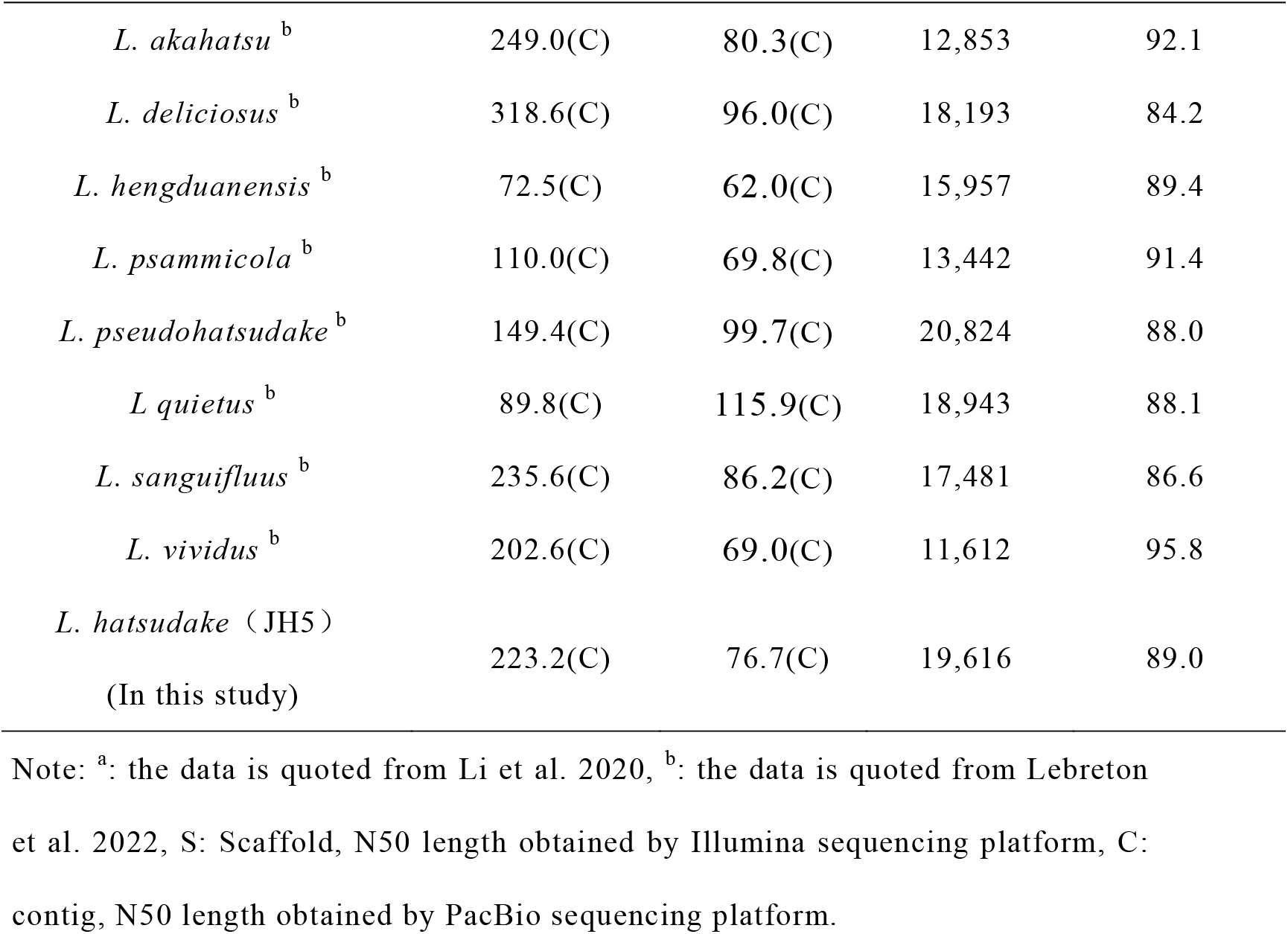
BUSCO evaluation comparison of *Lactarius* spp.

**Table 2.** Comparison of JH5, MG20 and 109 genome assembly and annotation

| Sample ID | JH5 | MG20 | 109 |
| --- | --- | --- | --- |
| Scaffolds | 312 | 36,963 | 815 |
| Contigs | 312 | 37,440 | 815 |
| Max Length of<br>contigs(bp) | 3,148,306 | 85,124 | - |
| N50 Length of contigs<br>(bp) | 223,180 | 5,268 | 261,250 |
| N90 Length of contigs<br>(bp) | 54,521 | 519 | - |
| Total length (Mb) | 76.7 | 73.8 | 95.5 |
| GC (%) | 54.4 | 51.9 | 52.1 |
| Gene number (#) | 19,616 | 18,513 | 16831 |
| Gene total length (bp) | 34,630,264 | - | - |
| Gene average length (bp) | 1,765 | - | - |
| Gene length/Genome (%) | 45.1 | - | - |

**Figure 1.**
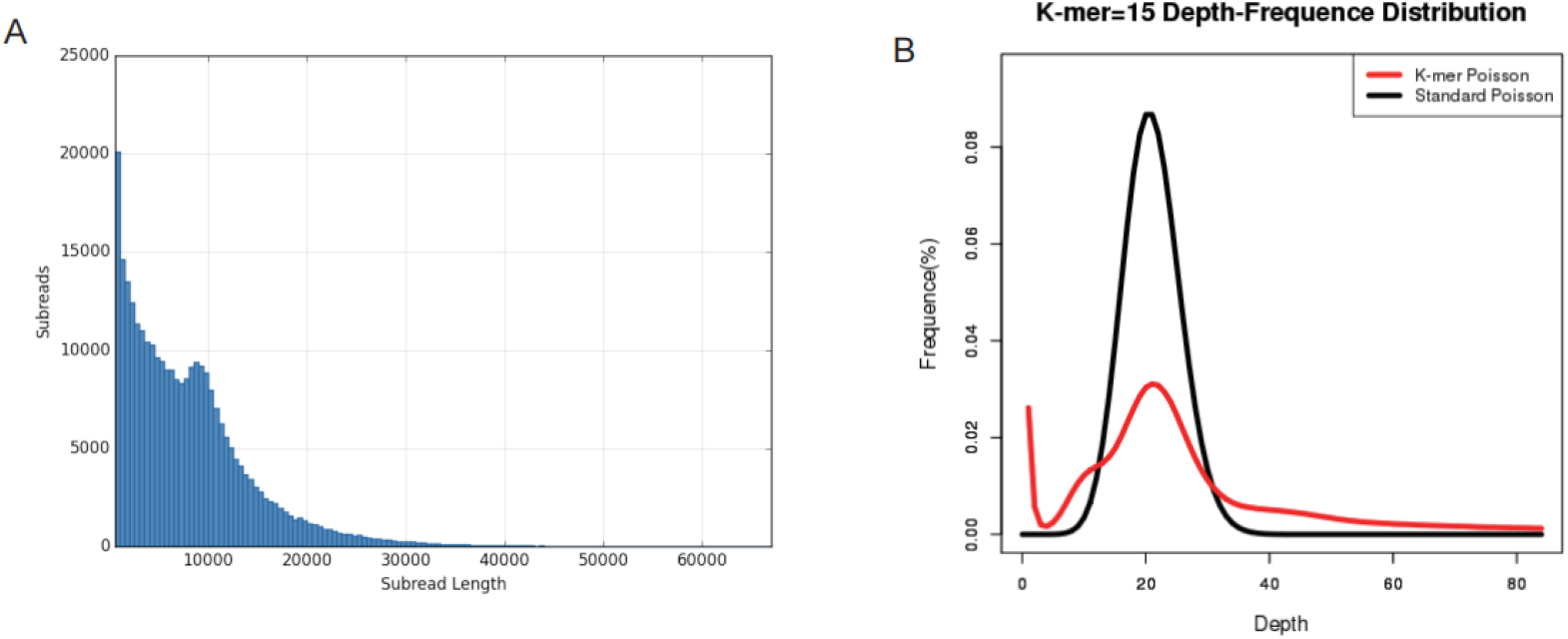
Read statistics and the distribution of Kmers. (A) The sequence read length distribution map of each sample library (the abscissa represents sequence read length, and the ordinate represents the number of reads corresponding to sequence read length). (B) Sample 15-mer statistical figure (the abscissa indicates the kmer depth, the ordinate indicates the proportion of the frequencies at each depth to the total number of the frequency. The red curve is the 15-mer depth distribution curve of the sequencing data, and the black curve is the standard Poisson distribution curve closest to it).

**Figure 2.**
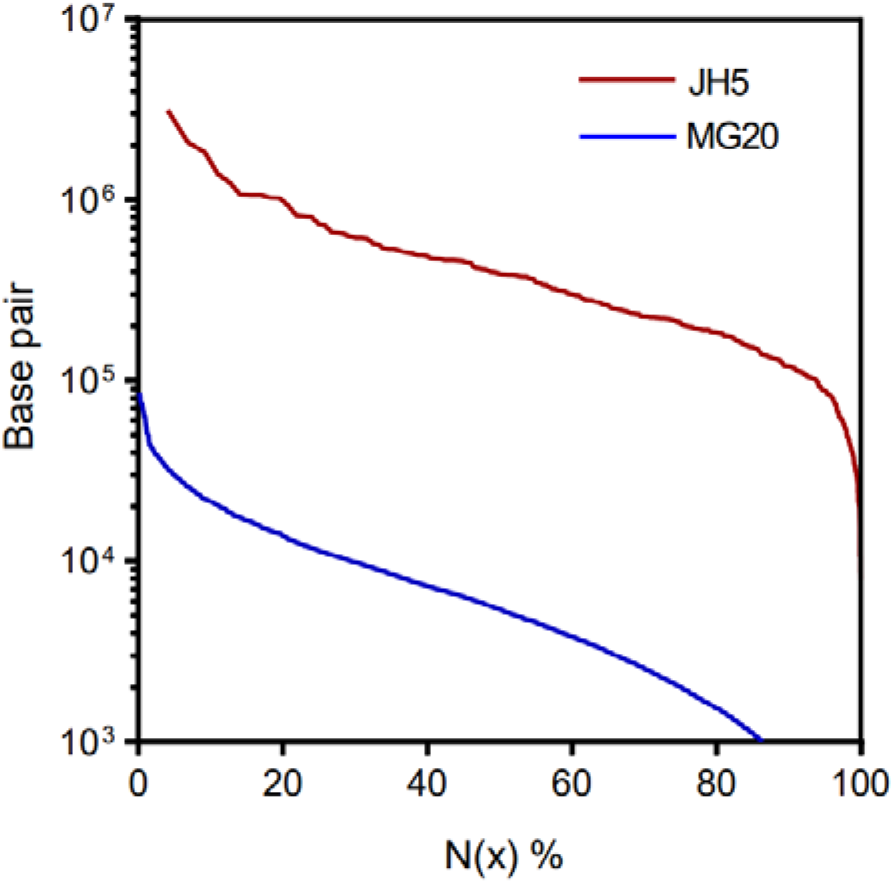
Comparison of length of Scaffold N(x) distribution between *L. hatsudake* JH5 and MG20

### Identification of repetitive sequences

In this study, a total of 33,787 repeat sequences were predicted within the *L. hatsudake* JH5 genome. Among these, the number of long terminal repeat sequences (LTR) was 12,436, which accounted for 10.18%, at an average length of 651 bp. This was followed by tandem repeat sequences (TR), which consisted of 10,801 sequence (1.57% of total bases), long interspersed nuclear elements (LINEs) (0.19%), short interspersed nuclear elements (SINEs) (0.13%), minisatellite DNA (0.57%), and microsatellite DNA (0.09%) (Table 3).

**Table 3.** Prediction of JH5 repeat sequences of samples

| Type | Number<br>(#) | Total length<br>(bp) | In genome<br>(%) | Average<br>length (bp) |
| --- | --- | --- | --- | --- |
| LTR | 12,436 | 7,806,839 | 10.1769 | 651 |
| DNA | 1,065 | 146,746 | 0.1913 | 142 |
| LINE | 912 | 98,384 | 0.1283 | 116 |
| SINE | 120 | 7,861 | 0.0102 | 67 |
| TR | 10,801 | 1,201,033 | 1.5656 | 112 |
| Minisa-ellite<br>DNA | 7,224 | 439,810 | 0.5733 | 61 |
| Microsatellite<br>DNA | 1,229 | 68,531 | 0.0893 | 56 |

Note: LTR: Long Terminal Repeat sequences; DNA: DNA transposon; LINE: Long interspersed nuclear elements; SINE: Short interspersed nuclear elements; TR: Tandem Repeat; Mini satellite DNA: microsatellite DNA; Microsatellite DNA: microsatellite.

### Identification of non-coding RNAs (ncRNA)

Non-coding RNAs (ncRNA) are a type of RNA that has been found to perform a variety of biological functions. It does not carry information that is translated into proteins though it still directly plays a role in activities at the RNA level (Bracher et al. 2020). Among microbes, sRNA, rRNA, and tRNA are the most commonly studied. For *L. hatsudake* JH5, tRNAs were found to be the most abundant, with a total length of 15.97 kb. This was followed by 5S rRNA, 18s and 28s, for a total of nine, with a total length of 8.0 kb. Additionally, there were 19 snRNAs, with an average length of 121 bp and a total length of 2.3 kb (Table 4).

**Table 4.** ncRNA statistical results after de-redundancy

| Type | Number (#) | Average length (bp) | Total length (kb) |
| --- | --- | --- | --- |
| tRNA | 204 | 78 | 16.0 |
| 5s (*de novo*) | 6 | 114 | 0.7 |
| 18s (*de novo*) | 2 | 1,644 | 3.3 |
| 28s (*de novo*) | 1 | 3,966 | 4.0 |
| snRNA | 19 | 121 | 2.3 |

Note: tRNA: Transfer RNA; snRNA: small nuclear RNA; rRNA: ribosomal RNA.

**Table 5.** Comparison of CAZy functional classification in *Lactarius* spp. and several other basidiomycetes

| Sample ID | CBM | CE | GH | GT | PL | AA | TOTAL |
| --- | --- | --- | --- | --- | --- | --- | --- |
| *L. hatsudake* (JH5) | 24 | 12 | 110 | 69 | 1 | 50 | 266 |
| *L. hatsudake* (MG20) | 12 | 10 | 27 | 12 | 2 | 9 | 72 |
| *L. deliciosus* (MG9) | 9 | 12 | 34 | 16 | 1 | 9 | 81 |
| *L. echinatus* (MG122) | 11 | 10 | 23 | 20 | 2 | 17 | 83 |
| *L. hygrophoroides* (MG19) | 9 | 9 | 24 | 16 | 0 | 14 | 72 |
| *L. indigo* (MG109) | 11 | 9 | 30 | 29 | 3 | 14 | 96 |
| *L. pinguis* (MG27) | 5 | 16 | 26 | 18 | 0 | 17 | 82 |
| *L. piperatus* (MG49) | 7 | 10 | 23 | 13 | 0 | 16 | 69 |
| *L. rugatus* (MG108) | 1 | 9 | 18 | 12 | 2 | 10 | 52 |
| *L. svolemus* (MG8) | 5 | 13 | 26 | 19 | 2 | 15 | 80 |
| *Boletus edulis* (MG6) | 12 | 16 | 34 | 13 | 2 | 14 | 91 |
| *T. calosporum* (MG102) | 9 | 13 | 22 | 19 | 1 | 13 | 77 |
| *L. edodes* | 58 | 31 | 245 | 75 | 9 | 85 | 461 |
| *P. chrysosporium* | 61 | 26 | 182 | 68 | 6 | 97 | 397 |
| *P. ostreatus* | 85 | 32 | 231 | 65 | 23 | 131 | 521 |
| *H. erinaceus* | 4 | 26 | 161 | 59 | 7 | 84 | 341 |

### Gene function analysis

The EVM pipeline was used to predict the PCGs of the *L. hatsudake* JH5 genome, with a total of 19,616 gene models identified and 14261 (72.70%) were expressed PCG models based on the transcriptome data. According to gene function analysis, all of the predicted genes were annotated using eleven databases (Figure 3A), including NR, KEGG, GO, etc. Among the NR databases, the KEGG, GO and Pfam databases produced 11,495 annotations (22.20%). The KEGG database annotated 10,403 (20.09%) while GO and Pfam results were the same at 11,538 annotations (22.28%). The SwissProt and KOG database annotated 2,265 and 1,794 genes, accounting for 4.37% and 3.46%, respectively. Lastly, the P450 and CAZy databases had the lowest number of genes annotated, accounting for 0.43% and 0.51%, respectively.

**Figure 3A:**
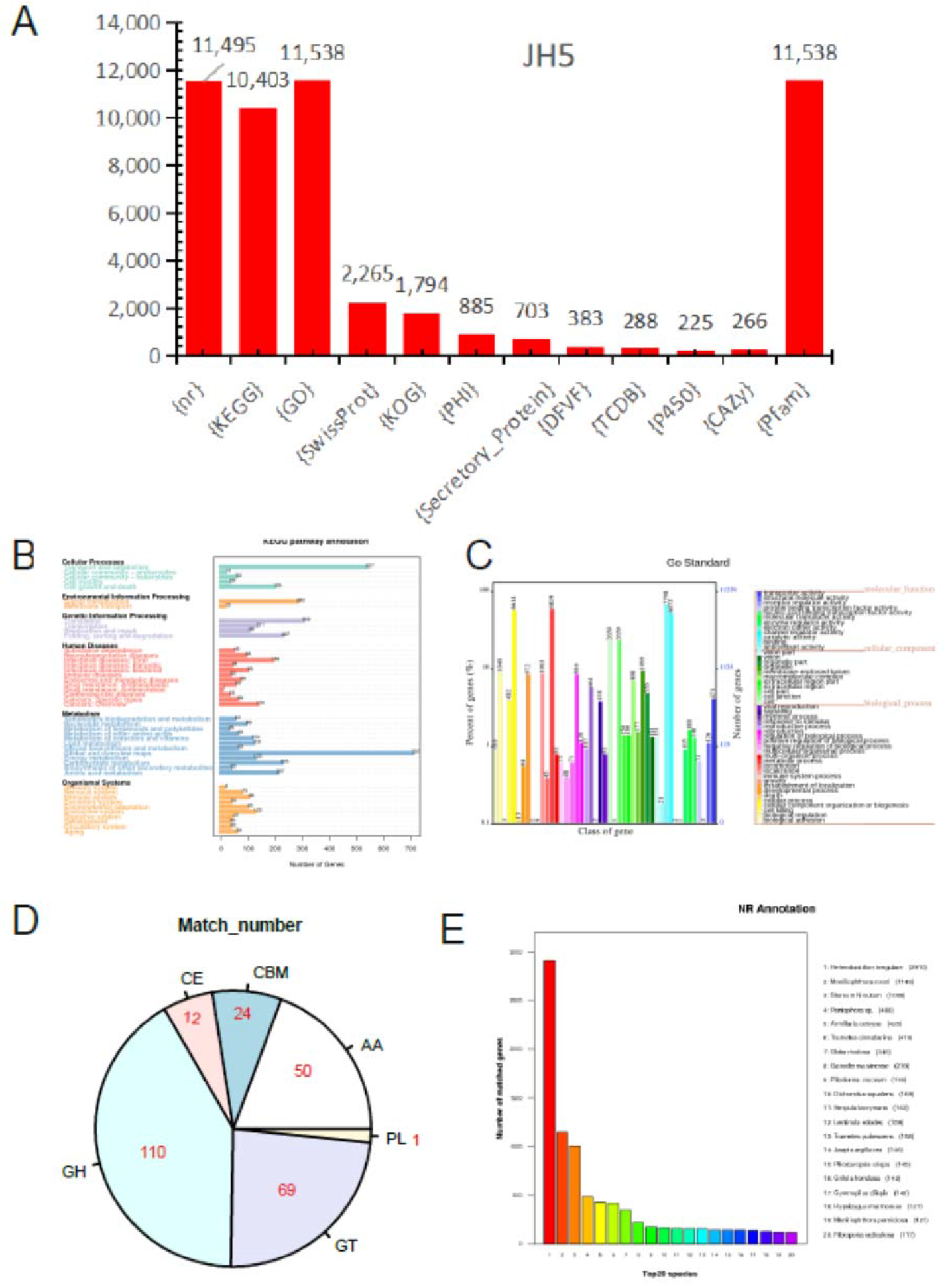
Statistical chart of the results of gene function analysis (the columns in the figure represent the number of genes annotated by the coding genes in each database). 3B: Gene functional annotation based on the GO functional classification map (the abscissa represents the GO functional classification on the sample annotation, the right ordinate indicates the number of genes on the annotation, and the left ordinate indicates the percentage of the number of genes in the annotation as a percentage of all encoded genes). 3C: Gene function annotation based on the KEGG metabolic pathway classification map (the number on the bar chart represents the number of genes on the annotation; the other axis is the code of each functional class of level1 in the database, the code is explained in the corresponding illustration). 3D: CAZy functional classification and corresponding gene quantity map. 3E: NR database species annotation statistical chart (Abscissa represents species ID, ordinate indicates the number of genes on the annotation).

### GO database annotation

The GO database was used to interpret annotated genes across three aspects: Cellular components, molecular functions, and biological processes. A total of 11,538 genes were annotated to the GO database, with cellular components accounting for 19.74%, molecular functions for 34.57%, and biological processes for 45.68%. Among these, the number of relationships to “binding” was the largest, reaching 7,798, followed by the metabolic process at 6,819, cell transformation at 6,614, and “catalytic activity” at 6,073 (Figure 3B).

### KEGG database annotation

The KEGG database was implemented in order to annotate genes covering the aspects of cellular process, environmental information processing, genetic Information processing, human diseases, metabolism and organismal systems. Metabolism accounted for the largest proportion (35.20%) while the number of genes annotated to transport and catabolism was maximum, followed by human diseases (16.20%) with the largest number of annotated genes annotated as “Viral” (Figure 3C).

### Carbohydrate enzymes database annotation

A database of carbohydrate enzymes was used that included a family of enzymes that can catalyze carbohydrate degradation, modification, and biosynthesis within five principal categories: glycoside hydrolases (GHs), glycosyl transferases (GTs), polysaccharide lyases (PLs), carbohydrate esterases (CEs), and auxiliary activities (AAs). The GH family comprised the largest proportion, with 110 annotated genes accounting for 41.35% (Figure 3D). The GT family annotated 69 genes that contained 25.94% but the PL family annotated only one family that comprised only 0.38%. The largest abundances within the GH family indicates that JH5 plays an important role in the formation of monosaccharides, oligosaccharides or carbohydrate complexes, the synthesis of alkyl glycosides and aromatic glycosides, the glycosylation of amino acids and peptides, and the glycosylation of antibiotics (Figure 3D). The comparison of CAZy functional classification in *Lactarius s*pp. showed that the annotated gene numbers of JH5 were higher than other *Lactarius s*pp. (Table 4). We also found that the number of carbohydrate enzymes of *L. hatsudake* JH5 was lower than saprophytic fungi such as *Lentinula edodes, Phanerochaete chrysosporium, Pleurotus ostreatus* and *Hericium erinaceus*. The results showed that perhaps due to the nature of JH5 as symbiotic mycorrhizal fungi, there were few CAZymes identified in *Lactarius*, with the capability of carbohydrate degradation weaker than that of white rot fungi. However, the numbers of GH and GT families in *L. hatsudake* JH5 were higher than other edible mycorrhizal fungi such as *L. hatsudake* MG20, other *Lactarius* spp., *Tuber calosporum* and *Boletus edulis*. As such, it appears that the capability of carbohydrate degradation is stronger than that of other mycorrhizal fungi of *Lactarius*.

### NR database annotation

The annotation of the NR database resulted in a total of 11,495 genes, of which JH5 exhibited the highest similarity with *Heterobasidion irregulare*. Here, 2,910 genes were annotated to the species for a total of 25.36% of all genes. This was followed by similarity to *Moniliophthora roreri* and *Stereum hirsutum*, with 1,148 (9.99%) and 1,008 (8.77%) genes shared, respectively *Fibroporia radiculosa*, with 117 genes. Of these most close genomic relatives, *H. irregulare* is known to cause Korean pine root rot (Blanchette et al. 2015), *S. hirsutum* is a crustal fungus that can secrete “red latex glue” (Kuo2008), *Obba rivulosa* can secrete “sap” (Kontro et al. 2020; Mainardi et al. 2018), and *Peniophora* sp. is capable of producing lactose. These data indicate that the majority of the genetic annotations of *L. hatsudake* strain JH5 are related to “mycorrhiza” and “red juice". Through comparison of the NR database, this genome was found to exhibit a high degree of similarity with other genomes, likely since the species are closely related and genes have not undergone major sequence differentiation (Figure 3E).

### Conclusion

In order to improve the genome assembly of *L. hatsudake*, we performed *de novo* sequencing and assembly of *L. hatsudake* JH5 by combining Illumina and PacBio sequencing. A total sequence length of 76,7 Mb of JH5 genome was assembled into 312 scaffolds with an N50 of 223.2 kb, and encoded 19,616 putative predicted genes. Compared with the released *Lactarius* spp. genomes, JH5 genome assembly presented the improved completeness and integrity. Here, the high-quality genome assembly of JH5 provides important insights into the biology of *L. hatsudake*. In addition, the identified genes may enhance our understanding of predicted gene function, enabling the study of biosynthesis of active compounds. Further research could focus on genes associated with growth and development or the biosynthesis of secondary metabolites. Although incomplete, the basic information provided by the elucidation of the *L. hatsudake* genome in this study is a novel attempt to facilitate biologically and agriculturally based research and thus support future applications of fungal species.

## Data availability

Sequencing data and genome assembly of JH5 for this project have been deposited in NCBI databases under project accession number PRJNA605941. Transcriptome data of JH5 were deposited into GenBank under the accession numbers of PRJNA841037.

## ACKNOWLEDGMENTS

This work was supported by the Hunan Key R&D Program (2016NK2161), Special Project of Forestry Science and technology innovation in Hunan Province(XLK201969, XLK201904), the National Key R&D Program of China (2017YFD060030203), Innovation Platform and talent plan of Hunan Province (2021NK4186).

## LITERATURE CITED

Blanchette RA, Held BW, Molloy D, Blake J, and D’Amato AW, 2015 First report of Heterobasidion irregulare causing root rot and mortality of red pines in minnesota. Plant Dis 99:1038. https://doi.org/10.1094/pdis-11-14-1232-pdn

Borodovsky M, and McIninch J, 1993 Genmark: Parallel gene recognition for both DNA strands. Comput Chem 17(2):123–133. https://doi.org/10.1016/0097-8485(93)85004-V

Bracher L, Ferro I, Pulido-Quetglas C, Ruepp MD, Johnson R et al., 2020 Human vtrna1-1 levels modulate signaling pathways and regulate apoptosis in human cancer cells. Biomolecules 10(4):614. https://doi.org/10.3390/biom10040614

Chin CS, Peluso P, Sedlazeck FJ, Nattestad M, Concepcion GT et al., 2016 Phased diploid genome assembly with single-molecule real-time sequencing. Nat Methods 13(12):1050–1054. https://doi.org/10.1038/nmeth.4035

Fang LZ, Shao HJ, Yang WQ, and Liu JK, 2006 Two new azulene pigments from the fruiting bodies of the basidiomycete Lactarius hatsudake. Helv Chim Acta 89(7):1463–1466. https://doi.org/10.1002/hlca.200690147

Gardner PP, Daub J, Tate JG, Nawrocki EP, Kolbe DL et al., 2009 Rfam: updates to the RNA families database. Nucleic Acids Res. 37: D136– D140. https://doi.org/10.1093/nar/gkn766

Gong WB, Wang YH, Xie CL, Zhou YJ, Zhu ZH et al., 2020 Whole genome sequence of an edible and medicinal mushroom, Hericium erinaceus (Basidiomycota, Fungi). Genomics 112: 2393–2399 https://doi.org/10.1016/j.ygeno.2020.01.011

Haas BJ, Salzberg SL, Zhu W, Pertea M, Allen JE et al., 2008 Automated eukaryotic gene structure annotation using evidencemodeler and the program to assemble spliced alignments. Genome Biol 9:R7. https://doi.org/10.1186/gb-2008-9-1-r7

Hall I R, 2011 The cultivation of edible mycorrhizal mushrooms in plantation forests. 6th International Workshop on Edible Mycorrhizal Mushrooms.

Kontro J, Maltari R, Mikkila J, Kahkonen M, Makela MR et al., 2020 Applicability of recombinant laccases from the white-rot fungus Obba rivulosa for mediator-promoted oxidation of biorefinery lignin at low pH. Front Bioeng and Biotech 8. https://doi.org/10.3389/fbioe.2020.604497

Koren S, Walenz BP, Berlin K, Miller JR, Bergman NH et al., 2017 Canu: Scalable and accurate long-read assembly via adaptive k-mer weighting and repeat separation. Genome Res 27(5):722–736. https://doi.org/10.1101/gr.215087.116

Lagesen K, Hallin P, Rødland EA, Staerfeldt HH, Rognes T, and Ussery DW, 2007 RNAmmer: consistent and rapid annotation of ribosomal RNA genes. Nucleic Acids Res., 35(9): 3100–3108. https://doi.org/10.1093/nar/gkm160

Lebreton A, Tang N, Kuo A, LaButti K, Andreopoulos W et al., 2022 Comparative genomics reveals a dynamic genome evolution in the ectomycorrhizal milk-cap (Lactarius) mushrooms. New Phytol. 235(1):306–319. https://doi.org/10.1111/nph.18143.

Li HY, Wu SR, Ma X, Chen W, Zhang J et al., 2020 The genome sequences of 90 mushrooms. Sci Rep 10(1):9982 https://doi.org/10.1038/s41598-018-28303-2

Love J, Palmer J, Stajich J, Esserand T, Kastman E et al., 2019 nextgenusfs/funannotate: funannotate v1.5.2. https://doi.org/10.5281/zenodo.2576527.

Lowe TM and Eddy S R, 1997 tRNAscan-SE: a program for improved detection of transfer RNA genes in genomic sequence. Nucleic Acids Res. 25(5): 955–964. https://doi.org/10.1093/NAR/25.5.955

Mainardi PH, Feitosa VA, de Paiva LBB, Bonugli-Santos RC, Squina FM et al., 2018 Laccase production in bioreactor scale under saline condition by the marine-derived basidiomycete Peniophora sp. CBMAI 1063. Fungal Biol 122(5):302–309. https://doi.org/10.1016/j.funbio.2018.01.009

Marcais G, and Kingsford C, 2011 A fast, lock-free approach for efficient parallel counting of occurrences of k-mers. Bioinformatics 27(6):764–770. https://doi.org/10.1093/bioinformatics/btr011

Martin F, Aerts A, Ahrén D, Brun A, Danchin EG et al., 2008 The genome of Laccaria bicolor provides insights into mycorrhizal symbiosis. Nature 452(7183):88–92. https://doi.org/10.1038/nature06556

Martin F, Kohler A, Murat C, Balestrini R, Coutinho PM et al., 2010 Perigord black truffle genome uncovers evolutionary origins and mechanisms of symbiosis. Nature 464(7291):1033–1038. https://doi.org/10.1038/nature08867

Martin F, and Selosse MA, 2008 The Laccaria genome: A symbiont blueprint decoded. New Phytol 180(2):296–310. https://doi.org/10.1111/j.1469-8137.2008.02613.x

Miyazawa M, Kawauchi Y, and Matsuda N, 2010 Character impact odorants from wild mushroom (Lactarius hatsudake) used in Japanese traditional food. Flavour and Frag J 25(4):197–201. https://doi.org/10.1002/ffj.1977

Saier MH, Reddy VS, Tamang DG, and Vastermark A, 2014 The transporter classification database. Nucleic Acids Res 42(D1): D251– D258. https://doi.org/10.1093/nar/gkt1097

Simao FA, Waterhouse RM, Ioannidis P, Kriventseva EV, and Zdobnov EM, 2015 BUSCO: Assessing genome assembly and annotation completeness with single-copy orthologs. Bioinformatics 31(19):3210–3212. https://doi.org/10.1093/bioinformatics/btv351

Stanke M, and Waack S, 2003 Gene prediction with a hidden Markov model and a new intron submodel. Bioinformatics 19: II215–II225. https://doi.org/10.1093/bioinformatics/btg1080

Tako M, Dobashi Y, Tamaki Y, Konishi T, Yamada M et al., 2012 Identification of rare 6-deoxy-d-altrose from an edible mushroom (Lactarius lividatus). Carbohydrate Res 350:25–30. https://doi.org/10.1016/j.carres.2011.12.016

Tan Y, Shen AR, Tan ZM, and Shen BM, 2021 Optimization of culture medium and strains for Lactarius hatsudake. Hunan Forestry Science and Technology 48(6): 1–80. https://doi.org/10.3969/j. issn 1003– 5710. 2021. 06. 001

Tan Z M, 2005 A study on the biological characters of Lactarius hatsudake Tanaka and its semi-artificial cultivation [D]. Changsha: Hunan Agricultural University. https://kns.cnki.net/KCMS/detail/detail.aspx?dbname=CDFD9908&filename=2006126271.nh

Tan Z M, Zhang Z G, Bu X Y, Fu S C, and Zhang P, 2006 Morphological and structural features of Lactarius hatsudake and L. hatsudake-Pinus massoniana ectomycorrhiza. Journal of Edible Fungi 13(2):36–44. https://doi.org/10.16488/j.cnki.1005-9873.2006.02.009

Tan Z M, Shen A R, and Fu S C, 2007 Successful cultivation of Lactarius hatsutake -an evaluation with molecular methods. Opera Mycologia (1): 38–41. https://doi.org/10.16488/j.cnki.1005-9873.2008.03.016

Vurture GW, Sedlazeck FJ, Nattestad M, Underwood CJ, Fang H et al., 2017 Genomescope: Fast reference-free genome profiling from short reads. Bioinformatics 33(14):2202–2204. https://doi.org/10.1093/bioinformatics/btx153

Walker BJ, Abeel T, Shea T, Priest M, Abouelliel A et al., 2014 Pilon: An integrated tool for comprehensive microbial variant detection and genome assembly improvement. Plos One 9(11). https://doi.org/10.1371/journal.pone.0112963

Wang Li, Li Zhuang, Zhu Mengjuan, Meng Li, Wang Hexiang et al., 2016 An acidic feruloyl esterase from the mushroom Lactarius hatsudake: A potential animal feed supplement. International Journal of Biological Macromolecules 93: 290–295. https://doi.org/10.1016/j.ijbiomac.2016.08.028

Zhang AL, Liu LP, Wang M, and Gao JM, 2007 Bioactive ergosterol derivatives isolated from the fungus Lactarius hatsudake. Chem Nat Compd 43(5):637–638. https://doi.org/10.1007/s10600-007-0215-x

